# Innate visual attraction in wood ants is a hardwired behavior seen across different motivational and ecological contexts

**DOI:** 10.1101/2021.01.29.428794

**Authors:** Cornelia Buehlmann, Paul Graham

## Abstract

Ants are expert navigators combining innate and learnt navigational strategies. Whereas we know that the ants’ feeding state segregates visual navigational memories in ants navigating along a learnt route, it is an open question if the motivational state also affects the ants’ innate visual preferences. Wood ant foragers show an innate attraction to conspicuous visual cues. These foragers inhabit cluttered woodland habitat and feed on honeydew from aphids on trees, hence, the attraction to ‘tree-like’ objects might be an ecologically relevant behavior that is tailored to the wood ants’ foraging ecology. Foragers from other ant species with different foraging ecologies show very different innate attractions. We investigated here the innate visual response of wood ant foragers with different motivational states, i.e., unfed or fed, as well as males that show no foraging activity. Our results show that ants from all three groups orient towards a prominent visual cue, i.e., this intrinsic visuomotor response is not context dependent, but a hardwired behavior seen across different motivational and ecological contexts.

## INTRODUCTION

Ants cleverly combine innate and learnt navigational strategies to travel between their nest and a feeding site (Buehlmann et al., 2020b; Knaden and Graham, 2016). Innate strategies such as path integration (Collett et al., 2001; Mueller and Wehner, 2010), pheromone trails (Harrison et al., 1989), or innately attractive visual cues (Collett, 2010; Graham et al., 2003) are key when unfamiliar with an environment. These innate responses can structure the ants’ paths, hence act as a scaffold and facilitate the learning of information relevant for navigation.

Conspicuous objects initiate innate behavior in many insects, including ants (fruit flies: (Gotz, 1987; Strauss and Pichler, 1998; Wehner, 1972), locusts: (Wallace, 1962), ladybirds: (Collett, 1988), mantids: (Poteser and Kral, 1995), leaf hoppers: (Brackenbury, 1996), ants: (Buehlmann et al., 2020a; Buehlmann et al., 2020c; Collett, 2010; Graham et al., 2003; Heusser and Wehner, 2002; Voss, 1967)), with many of these innate behaviors being ecologically tuned (e.g. prey detection, predator avoidance or landing site detection). Similarly, innate visual responses of different ant species are tailored to the ants’ habitat. Whereas wood ant foragers are attracted to large conspicuous objects (Buehlmann et al., 2020a; Buehlmann et al., 2020c; Graham et al., 2003; Voss, 1967), desert ants avoid them (Collett, 2010; Heusser and Wehner, 2002). Wood ants (*Formica rufa*) inhabit cluttered woodland habitat where they predominantly feed on honeydew from aphids on trees (Domisch et al., 2016). *Cataglyphis fortis* desert ants forage on dead arthropods that are unpredictably distributed on the ground in the food-scarce terrain of the Saharan salt pans (Buehlmann et al., 2014). *C. fortis* avoid large objects such as bushes, potentially to avoid predators. Hence, the differences in the innate bias of wood and desert ant foragers make sense from an ecological point of view and seem to be tuned to their foraging ecology. Furthermore, behavioral experiments in *Drosophila* have shown that both flying and walking fruit flies show innate visual responses tuned to the flies’ behavior requirements (Grabowska et al., 2018; Maimon et al., 2008), with small objects being aversive and large thin objects being attractive. Aversive behavior towards small objects potentially helps flies to avoid collision with other flying insects or predators, whereas bar-like objects may represent attractive feeding sites (Maimon et al., 2008).

Olfactory and visual information is commonly used to localize a food source, hence, the question arises if olfactory and visual responses vary with the feeding state. Indeed, behavioral experiments in fruit flies and parasitotic wasps have revealed that the animals’ feeding state can modulate innate olfactory and/or visual responses (Vogt et al., 2021; Waeckers, 1994). Experienced ant foragers can learn visual information for navigation (Graham and Philippides, 2017; Wehner et al., 2014; Zeil, 2012) and, we know that the ants’ motivational state (i.e. fed vs unfed) plays an important role in the use of learnt visual information (Harris et al., 2005; Wehner et al., 2006). More specifically, visual memories are primed by the ants’ feeding state and this controls the choice between foodward and homeward route memories (Harris et al., 2005). What we do not know, however, is whether the innate visual responses vary with an ants’ motivational state or caste.

To test this, we recorded the innate visual response of wood ant foragers with different motivational states, i.e., unfed or fed, as well as males that show no foraging activity (Hoelldobler and Wilson, 1990). We found that ants from all three groups orient towards the visual cue, i.e., the wood ants’ innate visual attraction is not flexible, but a hardwired behavior seen across sex and feeding state.

## METHODS

### Ants

Experiments were performed with laboratory kept wood ants *Formica rufa L*. collected in summer (between early August and mid September) from Ashdown forest, East Sussex, UK. Ants were kept in the laboratory under a 12 h light: 12 h darkness cycle at 25-27° C. Ants were fed *ad libitum* with sucrose (67% water and 33% sugar, by volume), dead crickets and water. Two days prior and throughout the experiments, sucrose and dead crickets in the nest were removed to increase the ants’ foraging motivation, but water was permanently available.

We recorded the innate visual response from naïve wood ants from three different groups: unfed female foragers, fed female foragers and unfed males. Unfed ants were taken from the nest and recorded in the behavioral arena. To get fed foragers, unfed foragers were taken from the nest and put into a 30 cm x 20 cm box where they were fed ad libitum with sucrose. Once they finished feeding, we left them in the feeding box for an extra 10 min before behavioral recordings to ensure that they did not have any motivation to return to the food. Our third experimental group were males, which are rarely spotted in lab kept wood ant colonies. Males were taken from the nest, and we only recorded and included data from those males that walked on the experimental platform. Males are winged, however, they are not good flyers, and approximately 80% of the released males walked.

Males were recorded in February 2020 from a nest that was collected in summer 2019. Fed and unfed foragers were recorded in October and November 2020 from a nest that has been in the lab for approximately three months. To test for nest effects, we included data from an additional group of unfed foragers that had been recorded in February 2018. These ants were taken from a nest that was collected in summer 2017, hence, the nest had been in the lab for five to six months at the time of the experiment. Ants from 2018 (n = 41 ants) and 2020 (n = 49 ants) approached the visual cue and their heading directions did not differ (Watson-Williams test, p > 0.05). In the absence of the visual cue, ants from both groups were not directed (2018: 11 ants; 2020: 21 ants; Rayleigh test, both p > 0.05). Hence, neither the nest nor the time spent in the lab had an influence on the ants’ heading direction and the data was thus pooled.

### Experimental setup

Ants from these three groups were released in the centre of a circular platform (120 cm in diameter) surrounded by a cylinder (diameter 3 m, height 1.8 m) with white walls (Fig. 1a). A 20° wide black rectangle (height: 90 cm, width: 52 cm) was placed on the inner wall of the surrounding cylinder. As a control, additional ants from the three groups were recorded when the visual cue was absent. Ants were only recorded once, and they had never been in this or any other behavioral setup before. Prior to this experiment ant colonies were kept in a featureless nest container. Ant’s perspective images from their nest and the experimental setup (with and without the visual cue) are shown in Supplementary Fig. 1. To decrease the directionality of potential olfactory traces, the surface of the platform was covered with white paper which was rotated between the tests of individual ants. Further, the visual cue was rotated between recordings to avoid the use of cues other than the black rectangle. The centre of the platform consisted of a cylindrical holding chamber of 6.5 cm diameter, which was remotely lowered to release the ant onto the platform. The ant’s position was recorded every 20 ms using a tracking video camera (Trackit, SciTrackS GmbH). The ants were recorded until they reached the edge of the platform.

**Fig. 1.**
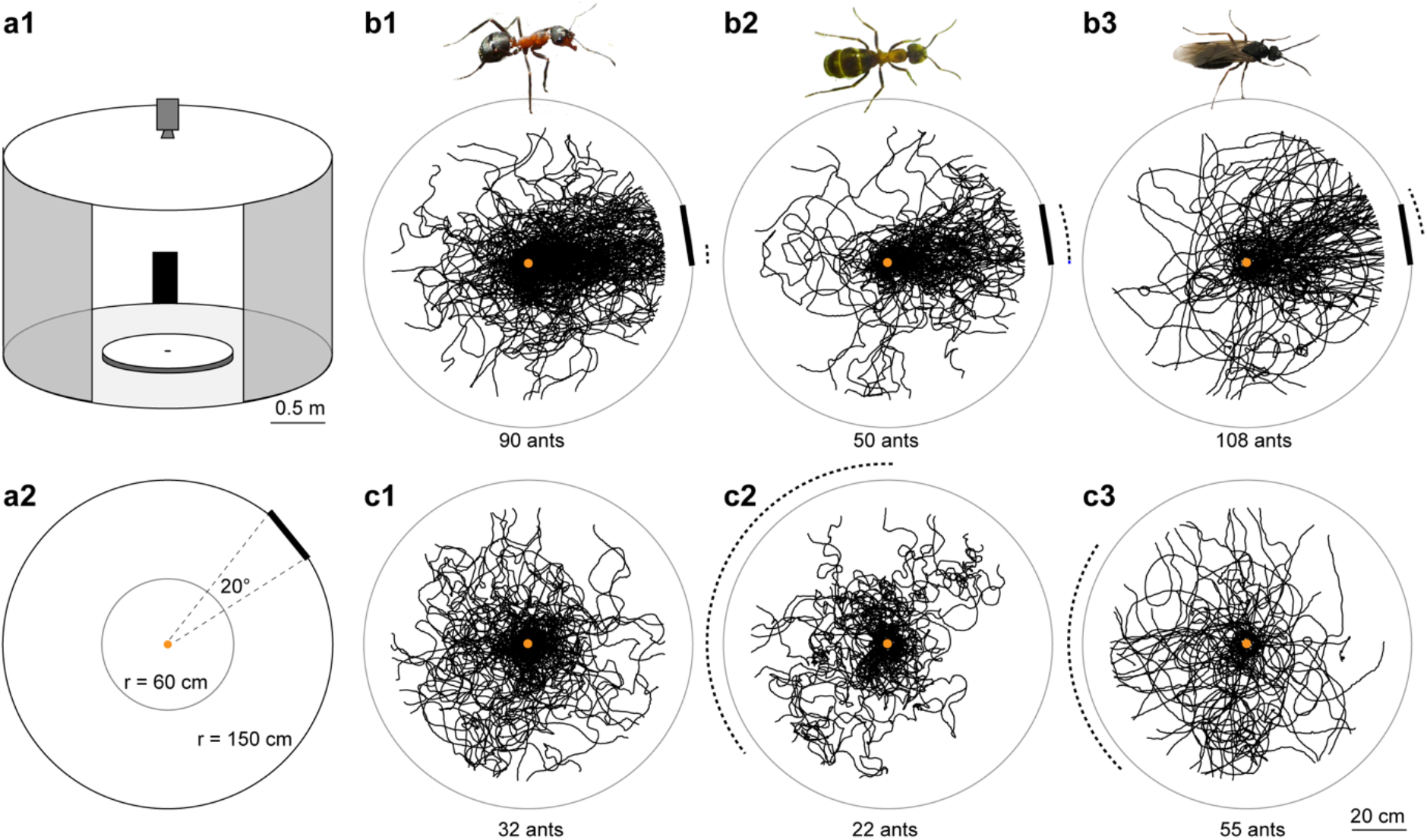
Innate visual attraction in wood ants is seen across different motivational and ecological contexts. **a1** Experimental arena in which naïve ants were recorded. Circular white platform (radius: 60 cm) is located in the centre of a cylinder (radius: 1.5 m, height: 1.8 m). A 20° wide black rectangle (height: 90 cm, width: 52 cm) is mounted at the inner wall of the surrounding cylinder. A camera recorded the ants’ paths from above. A small door permitted access to the arena shown here open and larger for clarity. **a2** A top-down view of the arena shown in a1. **b** Paths of ants released at the centre of the arena in the presence of the visual cue are shown as black lines. If the data is directed, dotted arcs show 95% confidence intervals (CIs) of the heading directions. The visual cue is shown at the platform edge instead of on the cylinder wall. b1: unfed foragers; b2: fed foragers; b3: unfed males. **c** As in b but ants were recorded without the visual cue. Sample sizes are shown in each panel. For statistics see also Supplementary Table 1.1. and 1.2.

### Analysis

Path analyses were done in MATLAB. Paths were trimmed at r = 50 cm and the ants’ final path directions were determined. Directionality of paths was tested using the Rayleigh test for circular data (Batschelet, 1981). Mean heading direction was considered to not deviate statistically from the visual cue direction if the visual cue laid fully or partially inside the 95% confidence interval (CI) of the ants’ heading direction. Different groups were compared using Watson Williams tests (Batschelet, 1981). Because of multiple pairwise testing, a Bonferroni correction was applied, and we used a corrected significance level of p = 0.0167. Additionally, average walking speed and path straightness (index of straightness = beeline distance / path length) were calculated for paths. A Kruskal Wallis test with Mann Whitney post-hoc tests and Bonferroni correction were used to compare ants from different groups.

## RESULTS

Paths from unfed foragers (n = 90 ants), fed foragers (n = 50 ants) and unfed males (n = 108 ants) were recorded in the behavioral arena in the presence of a conspicuous visual object (Fig. 1a). Ants from all three groups were directed (Rayleigh test; all p << 0.001; Fig. 1b; Supplementary Table 1.1). The 95% CIs for the ants’ final headings overlaid with the visual cue and the groups did not differ from each other (Watson Williams tests; all p > 0.0167, Bonferroni corrected p value; Supplementary Table 1.2). Whereas ants from all three groups approached the visual cue (Fig. 1b), we observed differences in the ants’ walking speed (Fig. 2a1, Supplementary Table 2) and path straightness (Fig. 2a2, Supplementary Table 2). Males walked significantly faster (Kruskal Wallis with Mann Whitney test and Bonferroni correction; unfed vs males, p < 0.05; fed vs males, p < 0.001) and straighter (Kruskal Wallis with Mann Whitney test and Bonferroni correction; unfed vs males, p < 0.001; fed vs males, p < 0.001) than fed and unfed foragers. Unfed and fed foragers differed in their walking speed (Kruskal Wallis with Mann Whitney test and Bonferroni correction; p < 0.001) but not in the path straightness (Kruskal Wallis with Mann Whitney test and Bonferroni correction; p > 0.05; Fig. 2).

**Fig. 2.**
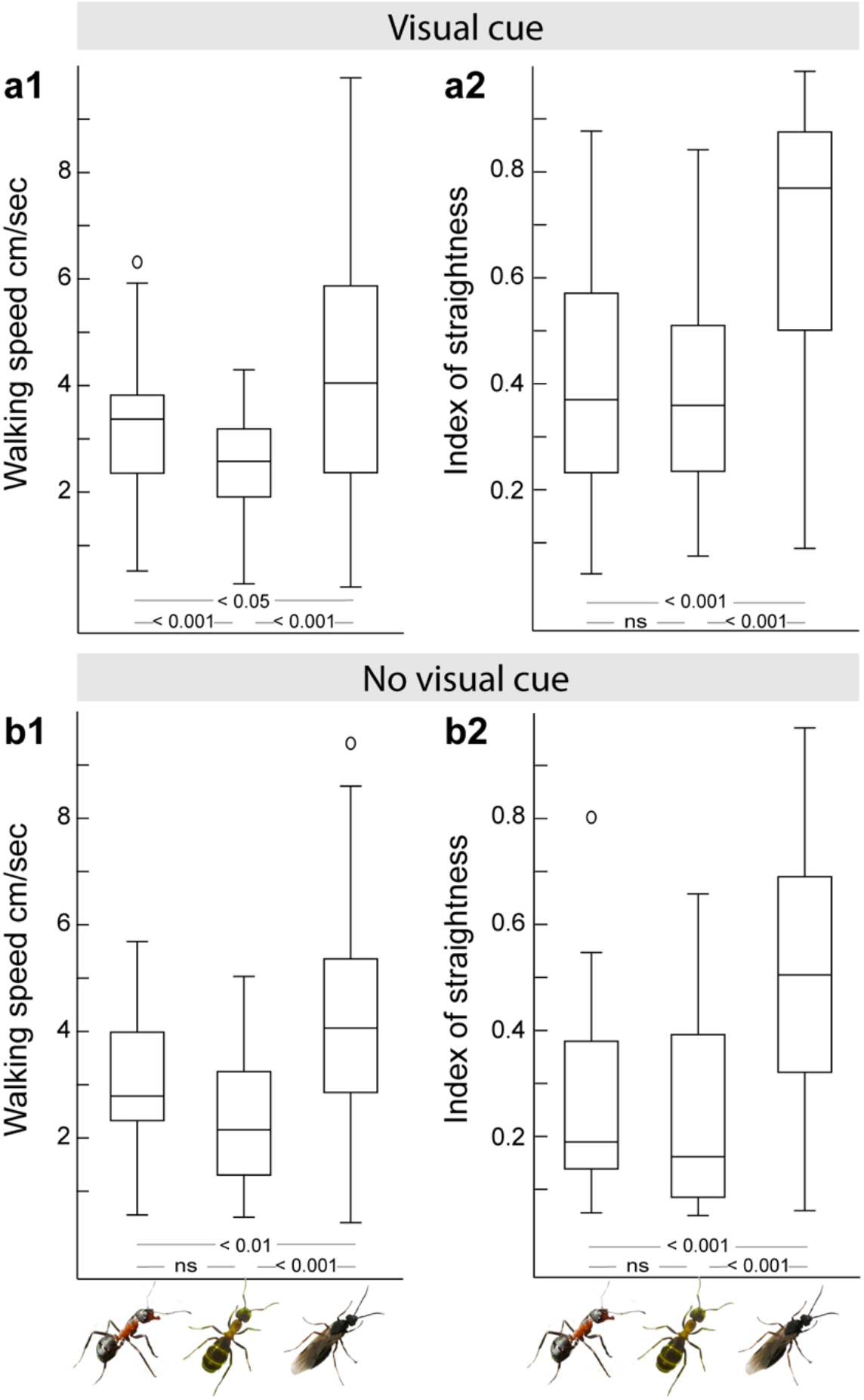
Walking speed and path straightness of ants with different motivational and ecological contexts. **a** Path details in the presence of the visual cue. Left, unfed, n = 90 ants; middle, fed, n = 50 ants; right, males, n = 108 ants. a1: Walking speed of ants differed significantly between the three groups. a2: Path straightness of males was significantly higher than of fed and unfed foragers, however, there was no difference between the paths of fed and unfed foragers. **b** Path details in the absence of the visual cue. Unfed, n = 32 ants; fed, n = 22 ants; males, n = 55 ants. b1: Walking speed of males was significantly higher than observed in unfed and fed foragers, but the latter two groups did not differ from each other. b2: Path straightness of males was significantly higher than of fed and unfed foragers but there was no difference between the paths of fed and unfed foragers. Boxplots: median, 25th and 75th percentiles (edges of the boxes) and whiskers for extreme values not considered as outliers (circles). Results of the Kruskal Wallis with Mann Whitney test and Bonferroni correction are shown in the figure. For statistics see also Supplementary Table 2.1. and 2.2.

In the absence of the visual cue (Fig. 1c), unfed foragers were not directed (Rayleigh test; p > 0.05, n = 32 ants), whereas the other two groups were directed (Rayleigh test; males, p < 0.01, n = 55 ants; fed foragers, p < 0.05, n = 22 ants) but showed a very large spread in directions (Supplementary Table 1.1). It is not obvious what the cause of this very small bias is (see Supplementary Fig. 1), possibly a small heterogeneity in the lighting or another aspect of the room. Importantly, in both the fed foragers and the males, these path headings differed significantly from the heading of the ants released in the presence of the visual cue (Watson-William test, both p << 0.001; Supplementary Table 1.2). Males walked significantly faster (Kruskal Wallis with Mann Whitney test and Bonferroni correction; unfed vs males, p < 0.01; fed vs males, p < 0.001) and straighter (Kruskal Wallis with Mann Whitney test and Bonferroni correction; unfed vs males, p < 0.001; fed vs males, p < 0.001) than fed and unfed foragers (Fig. 2b, Supplementary Table 2). Unfed and fed foragers did not differ from each other (Kruskal Wallis with Mann Whitney test and Bonferroni correction; unfed vs fed, both p > 0.05; Supplementary Table 2).

## DISCUSSION

We show here that the wood ants’ innate visual attraction is not context dependent, but an intrinsic visuomotor response seen across different motivational and ecological scenarios. More specifically, foragers with different motivational states, i.e., unfed or fed, as well as males that show no foraging behaviour naturally, all show an innate visual attraction to conspicuous visual objects (Fig. 1).

### Conspicuous objects initiate intrinsic visuomotor response

Many insects show intrinsic visuomotor response, i.e., they show fixed motor behaviors in response to specific visual input. Wood ant foragers show an innate attraction to large and conspicuous objects (Buehlmann et al., 2020a; Buehlmann et al., 2020c; Graham et al., 2003; Voss, 1967), whereas foragers from other ant species with different foraging ecologies show innate attractions different to wood ants (e.g. (Collett, 2010; Heusser and Wehner, 2002)). Wood ants inhabit cluttered woodland habitat and feed predominantly on honeydew from aphids on trees (Domisch et al., 2016), hence, the attraction to ‘tree-like’ objects might be an ecologically relevant behavior that is tailored to the wood ants’ foraging ecology (Graham and Wystrach, 2016).

Ants that rely on vision predominantly for navigation do not have a high visual resolution, but the ants’ wide-field and low-resolution vision allows robust visual navigation using the visual information provided by the environment. We do not know the exact details of the wood ants’ visual system, but the Australian desert ant that is a visual navigator in a similarly cluttered habitat has a horizontal visual field of approximately 150° per eye and an interommatidial angle of 3.7° (Schwarz et al., 2011). Which means that large conspicuous objects, such as tree trunks, would be easily perceivable (Supplementary Fig. 1).

### What is the role of innate behaviors in navigation?

We show here that innate visual orientation in wood ants is not controlled by the ants’ food related motivational state, however, innate behaviors so still play an important role during navigation for foraging. Path integration (e.g. (Collett et al., 2001; Mueller and Wehner, 2010)), pheromone trails (e.g. (Harrison et al., 1989)), or innately attractive visual cues (e.g. (Collett, 2010; Graham et al., 2003)) are innate strategies that are important for naive foragers or experienced ants exploring unfamiliar environments. As with path integration or odor trails, innate visual responses are intrinsic visuomotor responses that act as a scaffold by structuring the ants’ paths and thus facilitate the learning of sensory cues that are necessary for successful navigation (Graham et al., 2003). More specifically, incorporating these innate visual responses in route learning allows ants to take the same path repeatedly. Ants will thereby experience consistent views which accelerates the acquisition of visual information along the route. Additionally, it increases the robustness of learnt visual routes (Graham et al., 2003). Taken together, innate and learnt navigational strategies interact and allow the ants to successfully navigate between their nest and feeding sites (Buehlmann et al., 2020b; Knaden and Graham, 2016). In male ants it is possible that they are attracted to conspicuous objects to gain elevation to assist in dispersal. However, any role in innate visual orientation that assists in foraging would expect to be modulated by the ants’ motivational state (i.e. fed vs unfed foragers).

Experienced ant foragers navigate along routes using learnt visual information (Graham and Philippides, 2017; Wehner et al., 2014; Zeil, 2012). In a set of experiments, ant foragers were trained to learn visual cues along a route to a feeder and well-trained ants were subsequently tested with different motivational states (Harris et al., 2005; Wehner et al., 2006). Different behaviors were recorded in fed and unfed ant foragers, i.e., the ants’ motivational state played an important role in organizing visual navigational memories in ants navigating along learnt routes (Harris et al., 2005; Wehner et al., 2006). Hence, different to what we have shown here for innate visual responses, visual route memories are primed by the ants’ feeding state and this controls the choice between foodward and homeward route memories (Harris et al., 2005).

### Is it a surprise that innate visual responses are not flexible?

There are several examples of flexible innate behaviors in insects that can be modulated by the animals’ feeding state. For example, it was shown in parasitoid wasps that the insects’ individual feeding state controls their innate behavior (Waeckers, 1994). Unfed wasps are attracted by flower odors and yellow targets that indicate food while fed wasps are attracted to host odors and are not attracted by yellow colors (Waeckers, 1994). Other experiments have shown that the visual saliency can be modulated by the presence of odors. For instance, experiments in fruit flies have revealed that appetitive odors enhance the flies’ approach to visual cues (Cheng and Frye, 2021). Furthermore, experiments with hawkmoths have shown that the moths’ innate color preference depends on ambient light conditions (Kuenzinger et al., 2019). These moths are crepuscular, and their color preferences are tuned to illuminance and background. This flexible behavior allows them to successfully forage under different light conditions. These examples show that insects are equipped with innate visual preferences, but in some species necessary behavioral flexibility is maintained and behaviors are tuned to foraging ecology.

Given that there are many examples of flexibility, there is the question as to why wood ants do not show any flexibility in their innate visual orientation to conspicuous objects. One reason could be that ant foragers rely on their multimodal navigation toolkit which overcomes potential problems with an inflexible innate visual reflex. Fed foragers will have path integration information and learnt odor cues and visual information (Buehlmann et al., 2020b) as they attempt to return to their nest and these sources of orientation information may be more than enough to overcome innate visual attractions (Buehlmann et al., 2020c; Harris et al., 2007; Schwarz & Cheng, 2010) that may otherwise disrupt a homeward journey.

### Innate visual attraction in the insect brain

Recent research on the central complex (CX), a brain area located at the midline of the insect brain, revealed its importance in navigational tasks (Fisher, 2022). Work on ring neurons in the central complex of the fruit fly showed that many of these neurons preferentially respond to vertically oriented objects (Seelig and Jayaraman, 2013) and computational simulations further revealed that the sparse encoding in a population of these visually responsive ring neurons is sufficient to explain the innate behaviour observed in flies (Dewar et al., 2017; Wystrach et al., 2014). More specifically, the activity of these neurons was highest when the agent was facing a narrow vertical bar or an inner edge of a wide vertical bar. Furthermore, when exposed to natural scenes, these cells preferentially respond when facing large high contrast objects such as trees (Dewar et al., 2017; Wystrach et al., 2014). Finally, as previously mentioned, innate visual responses are intrinsic visuomotor responses that act as a scaffold by structuring paths and thus facilitate the learning of sensory cues that are necessary for navigation along a route (Graham et al., 2003). During this learning, innate behaviours are over-ridden but not changed, hence, the insect keeps the innate responses as a backup (Goulard et al., 2021). It thus makes sense that the ants’ motivational state controls visual memory along a learnt route (Harris et al., 2005; Wehner et al., 2006), but that there is no flexibility in intrinsic visuomotor responses (Fig. 1).

## DECLARATIONS

### Availability of data and material

The dataset generated during this study is available from the University of Sussex research repository (hosted by Figshare): https://doi.org/10.25377/sussex.14270330.

### Code availability

Not applicable.

### Ethics approval

Ethical approvals are not required by UK law for ant behavior experiments, but local permissions were still acquired from Sussex University.

### Consent to participate

Not applicable.

### Consent for publication

Not applicable.

