## Supplementary Fig. 1 for "Innate visual attraction in wood ants is a hardwired behavior seen across different motivational and ecological contexts"

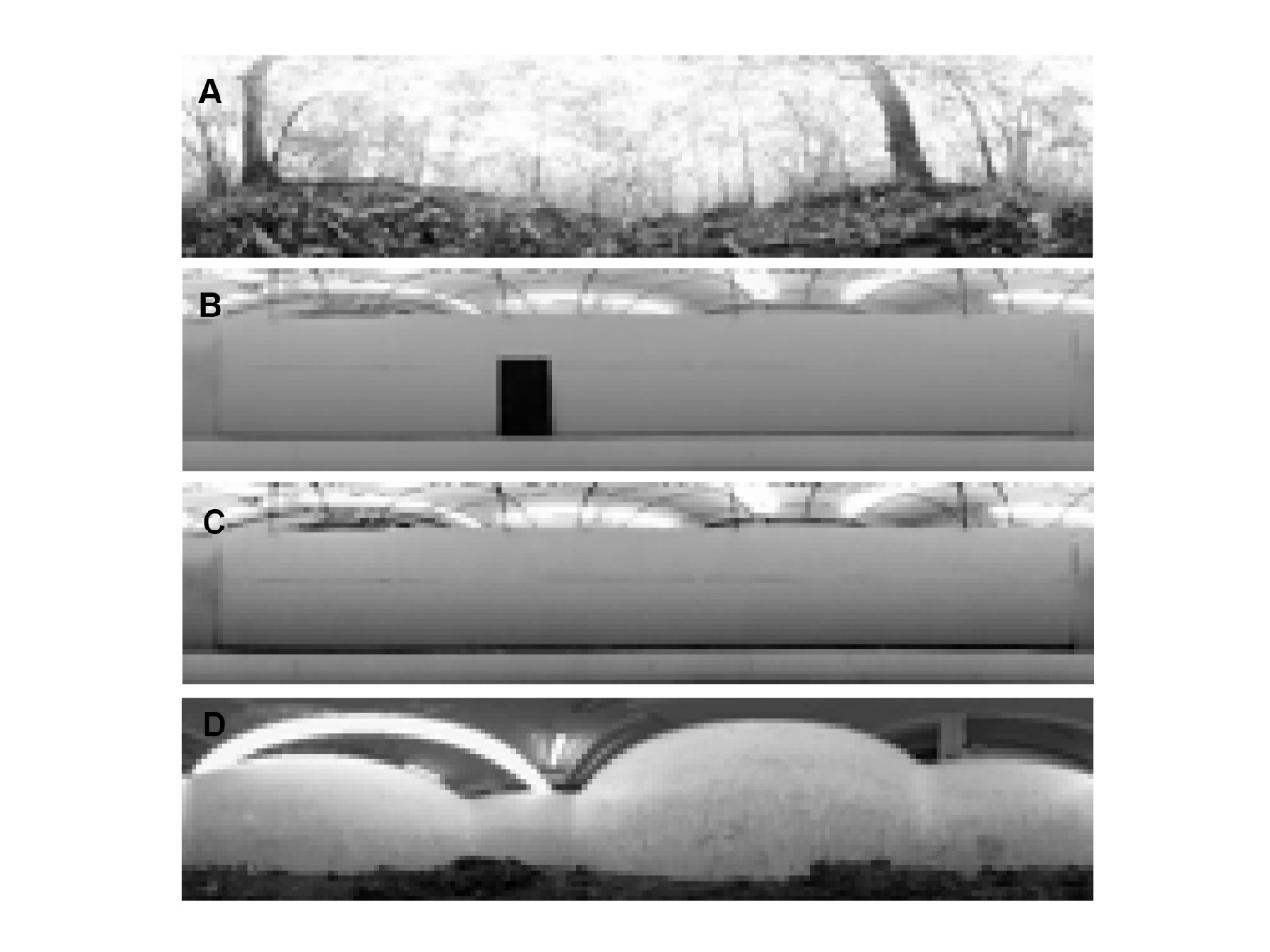

**Supplementary Fig 1.** Panoramic “ant’s eye” pictures from a natural wood-ant habitat in East Sussex (A), experimental setup with the visual cue (B), visual setup with no visual cue (C) and lab nest environment (D). Panoramic images were taken with a Kodak Pixpro SP360 4K and processed in Matlab. 360° view is shown with 4° resolution.
