## Supplementary Table 1 for "Innate visual attraction in wood ants is a hardwired behavior seen across different motivational and ecological contexts"

**Supplementary Table 1.1.** Heading directions

|  | Conditions | Sample size | Mean (°) | Confidence interval | Rayleigh test (p value) |
| --- | --- | --- | --- | --- | --- |
| Visual cue | Foragers unfed | 90 ants | 0.6 | 354.4 / 6.731 | << 0.001 |
|  | Foragers fed | 50 ants | 350.2 | 340.6 / 359.7 | << 0.001 |
|  | Males | 108 ants | 343.3 | 335.8 / 350.8 | << 0.001 |
| No visual cue | Foragers unfed | 32 ants | n/a | n/a | 0.27 |
|  | Foragers fed | 22 ants | 206.6 | 141 / 272.2 | 0.0468 |
|  | Males | 55 ants | 174.5 | 134.9 /214.0 | 0.0015 |

**Supplementary Table 1.2.** Heading directions: Comparisons

|  | Pairwise comparisons | Watson Williams tests (p value) |
| --- | --- | --- |
| Visual cue | Foragers unfed vs Foragers fed | 0.237 (a) |
|  | Foragers unfed vs Males | 0.021 (a) |
|  | Foragers fed vs Males | 0.463 (a) |
| No visual cue | Foragers fed no visual cue vs Foragers fed visual cue | << 0.001 |
|  | Males no visual cue vs Males visual cue | << 0.001 |

(a) Bonferroni corrected significance level of p = 0.0167.
