## Supplementary Table 2 for "Innate visual attraction in wood ants is a hardwired behavior seen across different motivational and ecological contexts"

**Supplementary Table 2.1.** Walking speed and path straightness

|  | Conditions | Sample size | Median  (Walking speed, cm/sec) | Median  (Path straightness) |
| --- | --- | --- | --- | --- |
| Visual cue | Foragers unfed | 90 ants | 3.3 | 0.37 |
|  | Foragers fed | 50 ants | 2.5 | 0.36 |
|  | Males | 108 ants | 4.0 | 0.77 |
| No visual cue | Foragers unfed | 32 ants | 2.8 | 0.19 |
|  | Foragers fed | 22 ants | 2.1 | 0.16 |
|  | Males | 55 ants | 4.1 | 0.51 |

**Supplementary Table 2.2.** Walking speed and path straightness: Comparisons

|  | Pairwise comparisons | Walking speed,  Kruskal Wallis with Mann Whitney test and Bonferroni correction (p value) | Path straightness  Kruskal Wallis with Mann Whitney test and Bonferroni correction (p value) |
| --- | --- | --- | --- |
| Visual cue | Foragers unfed vs Foragers fed | < 0.001 | 1 |
|  | Foragers unfed vs Males | 0.02 | < 0.001 |
|  | Foragers fed vs Males | < 0.001 | < 0.001 |
| No visual cue | Foragers unfed vs Foragers fed | 0.38 | 1 |
|  | Foragers unfed vs Males | 0.003 | < 0.001 |
|  | Foragers fed vs Males | <0.001 | < 0.001 |
